# Statistical modelling of CDR3 sequences provides robust quality control for TCR repertoire datasets

**DOI:** 10.64898/2025.12.11.693618

**Authors:** Dana Léa Moreno, Giancarlo Croce, David Gfeller

## Abstract

T-Cell Receptors (TCRs) show extensive sequence diversity across T cells. This diversity arises from different choices of V and J genes and from insertions and deletions at the V(D)J junction within the Complementary-Determining Region 3 (CDR3) loop. Here, we quantify how V and J gene usage shapes CDR3 length and amino acid composition. In repertoires of TCRs with either unknown or known specificity, we show that CDR3 length is strongly influenced by the number of germline-encoded CDR3 residues in V and J genes, and that, on average, 80% of CDR3α and 65% of CDR3β residues are determined by V and J gene usage. We further show that inconsistencies between V and J gene annotations and CDR3 sequences can be leveraged to identify potential issues in multiple TCR repertoire datasets. Overall, our study quantifies the impact of V and J gene usage on CDR3 length and amino acid composition and provides a robust framework for quality control of TCR repertoire sequencing data.

## Introduction

T cells are a fundamental component of the adaptive immune system. They are responsible for identifying infected or malignant cells, and initiating or coordinating immune responses. T-cell activation is elicited by the binding of the T-cell receptor (TCR) to antigen-derived peptides presented on major histocompatibility complex (MHC) molecules. The set of TCRs expressed by T cells in an individual is referred to as the TCR repertoire. TCR repertoires are characterized by a high diversity of TCR sequences, which ensures an effective immune response against a wide range of epitopes (i.e. peptide-MHC) (Dupic et al., 2019; Mora and Walczak, 2016; Nikolich-Žugich et al., 2004; Qi et al., 2014; Zarnitsyna et al., 2013). TCR repertoires encode a footprint of the past and current immunological status of individuals. For this reason, many studies have attempted to leverage TCR repertoires to support disease diagnostics, TCR-based biomarker discovery and TCR T-cell therapy (Bai et al., 2022; Charles et al., 2020; Dong et al., 2021; Greissl et al., 2022; Li et al., 2025; Minervina et al., 2021; Mitchell et al., 2023; Rosenberg et al., 2011; Zheng et al., 2021).

TCRs are heterodimers composed of an α and a β chain, with the α chain encoded at the TRA locus and the β chain at the TRB locus. During T cell development, germline-encoded genes for each chain are assembled through a stochastic process called V(D)J recombination. For the α chain, one variable (V) and one joining (J) gene are selected from a pool of germline-encoded genes and recombined. For the β chain, one short diversity (D) gene is additionally incorporated between the V and J. At the junction between the V (D) and J genes, random insertions, deletions and mutations of nucleotides occur. This highly variable region falls within the Complementarity-Determining Region 3 (CDR3), which encompasses the last part of the germline V gene, the junctional region, and the first part of the J gene. From a structural point of view, CDR3s are known to mediate many contacts with epitopes. For this reason, CDR3s have often been considered as the key region for TCR-epitope recognition and have been the primary focus of several studies (Beshnova et al., 2020; Meynard-Piganeau et al., 2024; Moris et al., 2021; Pham et al., 2023; Valkiers et al., 2021; Xu et al., 2023). The influence of V and J genes on CDR3 length and amino acid composition has been investigated in some studies (Freeman et al., 2009; Levi and Louzoun, 2022; Ma et al., 2016; Manfras et al., 1999; Nishio et al., 2004; Pannetier et al., 1993). For example, it is well established that the beginning and end of the CDR3 reflect the contributions of the V and J genes, respectively. However, a systematic assessment of how V and J genes shape CDR3 sequences across all V and J genes and all CDR3 lengths for both chains has not been performed, and many studies analyzing TCR repertoires treat CDR3 sequences independently of the V and J genes (Beshnova et al., 2020; Jiang et al., 2023; Meynard-Piganeau et al., 2024; Moris et al., 2021; Pham et al., 2023; Valkiers et al., 2021).

Databases such as iReceptor (Corrie et al., 2018) have collected TCR repertoire data from multiple studies, providing access to millions of TCRα and TCRβ sequences. TCR repertoires can be experimentally profiled using targeted amplicon sequencing that amplifies V and J gene regions with multiplex primer sets or 5′-RACE designs (Genolet et al., 2023; Heather et al., 2016; Mamedov et al., 2013; Robins et al., 2009). Many protocols incorporate unique molecular identifiers (UMIs) to mitigate PCR and sequencing errors and to improve quantitative accuracy (Shugay et al., 2014). After sequencing, pipelines such as MiXCR perform V(D)J assignment, CDR3 reconstruction, and clonotype assembly against reference databases (Alamyar et al., 2012; Bolotin et al., 2015; Ye et al., 2013). The most widely used reference database of germline V and J genes is the ImMunoGeneTics (IMGT) database (Giudicelli, 2004). These workflows produce annotated repertoires (i.e., V and J genes and CDR3 sequence for each TCR) that serve as the basis for the analysis of TCR repertoires across patients (Heather et al., 2017).

TCR sequencing data are prone to multiple sources of error that can significantly impact the quality of a dataset (Bolotin et al., 2012; Bradley and Thomas, 2019; Heather et al., 2017; Shugay et al., 2014). PCR and sequencing errors, as well as inaccuracies during the reconstruction of TCR sequences, can lead to the misidentification of V and J genes or inaccurate CDR3 sequences (Bolotin et al., 2012; Heather et al., 2017). In addition to these technical issues, multiple definitions of the CDR3 boundaries have been used across different studies, differing for instance in whether the conserved C at the beginning and the conserved F/W at the end of the CDR3 are included in the CDR3 definition. Diverse V and J gene nomenclatures have also been used (e.g., TRAV12-2 versus TCRAV12-02).

Some approaches have been developed to address these issues and improve the quality and usability of existing TCR repertoire datasets. For example, TidyTcells (Nagano and Chain, 2023) standardizes gene names to IMGT nomenclature and applies simple rules to harmonize CDR3 sequences by adding a cysteine (C) at the beginning of the CDR3 if this amino acid is missing and a phenylalanine (F) at the end of the CDR3 if F or W are missing. VDJtools (Shugay et al., 2015) provides a unified post-analysis framework across multiple input formats, including modules for basic repertoire statistics and V and J gene usage, diversity analysis, and repertoire clustering. Efforts to standardize adaptive immune receptor repertoire datasets (https://docs.airr-community.org) play a key role in addressing some of these issues.

In this study, we leverage the wealth of available TCR sequencing data to systematically examine how V and J gene usage shapes CDR3 length and amino acid composition. Building on these results, we introduce a robust quality-control framework that detects inconsistencies between V/J gene assignments and CDR3 amino acid sequences in TCR repertoire datasets.

## Results

### V and J gene usage shapes the length of CDR3 sequences

To systematically investigate the relationship between V and J gene choices and CDR3 length and amino acid composition, we collected TCR repertoire sequencing data from the iReceptor platform (Corrie et al., 2018). Samples from individuals under special immune conditions (e.g., leukemia) or demonstrating major biases in V and J usage or CDR3 length distributions were not included. After initial preprocessing steps (see Methods), our dataset contained 6,306,690 TCRα sequences from 25 subjects (Minervina et al., 2021; Rubelt et al., 2016; Shomuradova et al., 2020) and 145,357,767 TCRβ sequences from 833 subjects (DeWitt et al., 2015; Hannan et al., 2021; Jia et al., 2018; Joseph et al., 2022; Minervina et al., 2021; Nolan et al., 2025; Rubelt et al., 2016; Shomuradova et al., 2020) (Figure 1A and Supp. Table 1).

**Figure 1:**
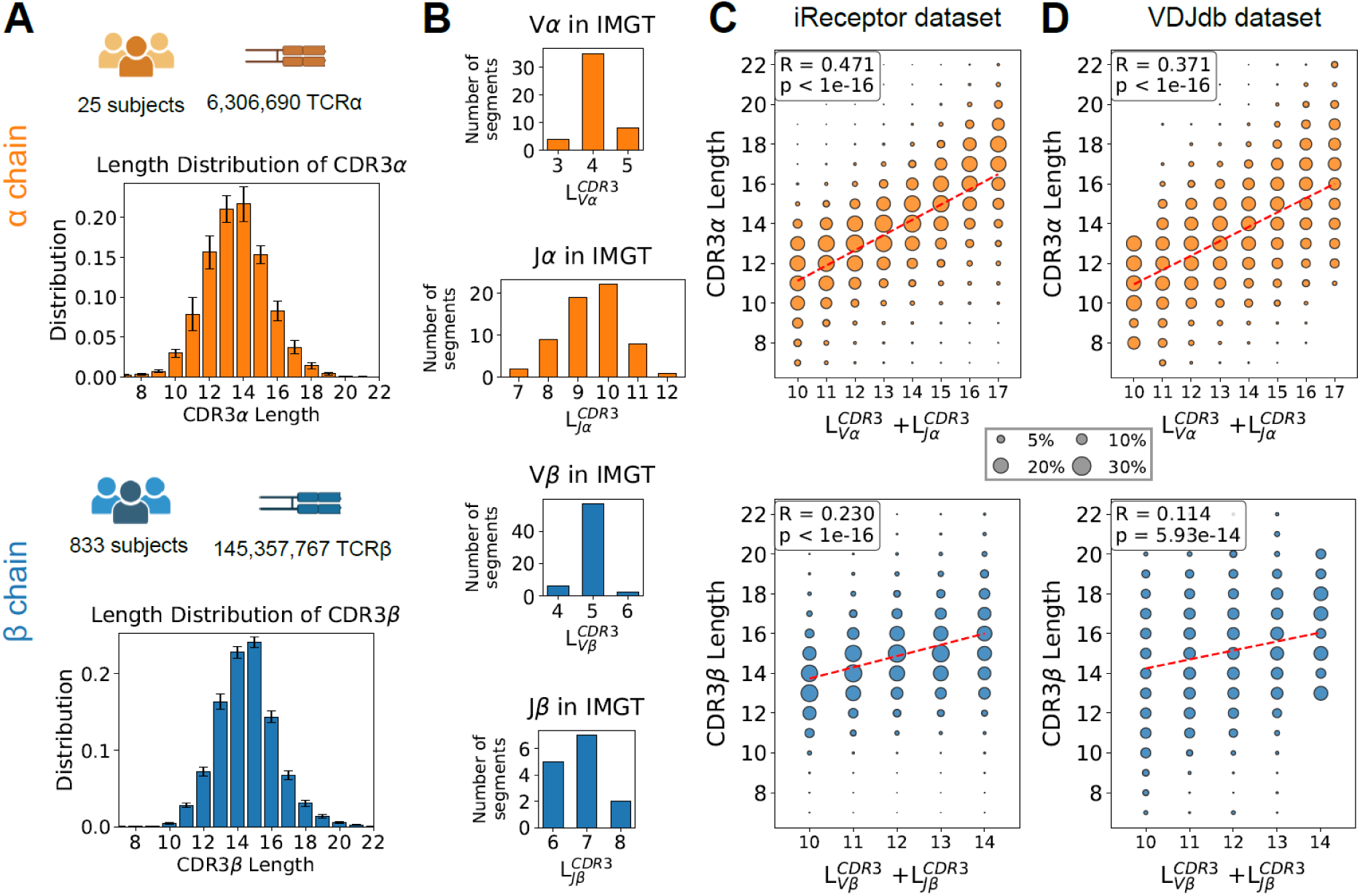
V and J gene usage shapes the length of CDR3 sequences. **(A)** Summary of the TCR datasets collected from the iReceptor database and distribution of CDR3 lengths for α and β chains. Error bars show the standard deviation across subjects. **(B)** Distribution of the number of CDR3 residues in germline V and J genes (L^CDR3^_v/j_) for α and β chains based on the IMGT reference database. **(C)** Correlations between the total number of CDR3 residues in germline V and J genes (L^CDR3^_v_ + L^CDR3^_j_) and the CDR3 lengths for α and β chains in repertoires of TCRs with undetermined specificities (iReceptor dataset). The size of each dot is proportional to the number of sequences. **(D)** Correlations between the total number of CDR3 residues in germline V and J genes (L^CDR3^_v_ + L^CDR3^_j_) and the CDR3 lengths for α and β chains in repertoires of TCRs with known specificities (VDJdb dataset). The size of each dot is proportional to the number of sequences.

We first examined the distribution of CDR3 sequence lengths in α and β chains. As observed in previous studies (Freeman et al., 2009; Liu et al., 2014), CDR3 lengths followed bell-shaped distributions, with maxima at 14 amino acids for the α chain and 15 amino acids for the β chain (Figure 1A). These distributions were consistent across all subjects analyzed in this work (cf. error bars in Figure 1A).

We next retrieved from IMGT the sequences of all germline-encoded V and J genes and extracted the number of CDR3 residues in germline V and J gene sequences (referred to as L^CDR3^_va_, L^CDR3^_vβ_, L^CDR3^_ja_ and L^CDR3^_jβ_). For V genes, L^CDR3^_v_ corresponds to the number of N-terminal residues of the V gene starting at the conserved cysteine (C). For J genes, L^CDR3^_j_ corresponds to the number of C-terminal residues of the J gene ending at the residue (i.e., F or W) before the conserved GXG motif. Figure 1B shows the distributions of L^CDR3^_v/j_ values for both chains, with the highest variability observed for Jα genes.

We then investigated for both chains the relationship between the total number of CDR3 residues in germline V and J genes (L^CDR3^_v_ + L^CDR3^_j_) and the length of CDR3 sequences across our collection of TCR repertoires. As shown in Figure 1C, we observed a significant correlation, particularly for the α chain. The individual contributions of specific gene families are provided in Supp. Figure 1A. The same observation could be made for TCRs with known specificity from the VDJdb database (Shugay et al., 2018) (Figure 1D and Supp. Figure 1B). Similar results were also obtained when analyzing murine TCR repertoire sequencing datasets (see Methods, Supp. Figure 2 and Supp. Table 2).

**Figure 2:**
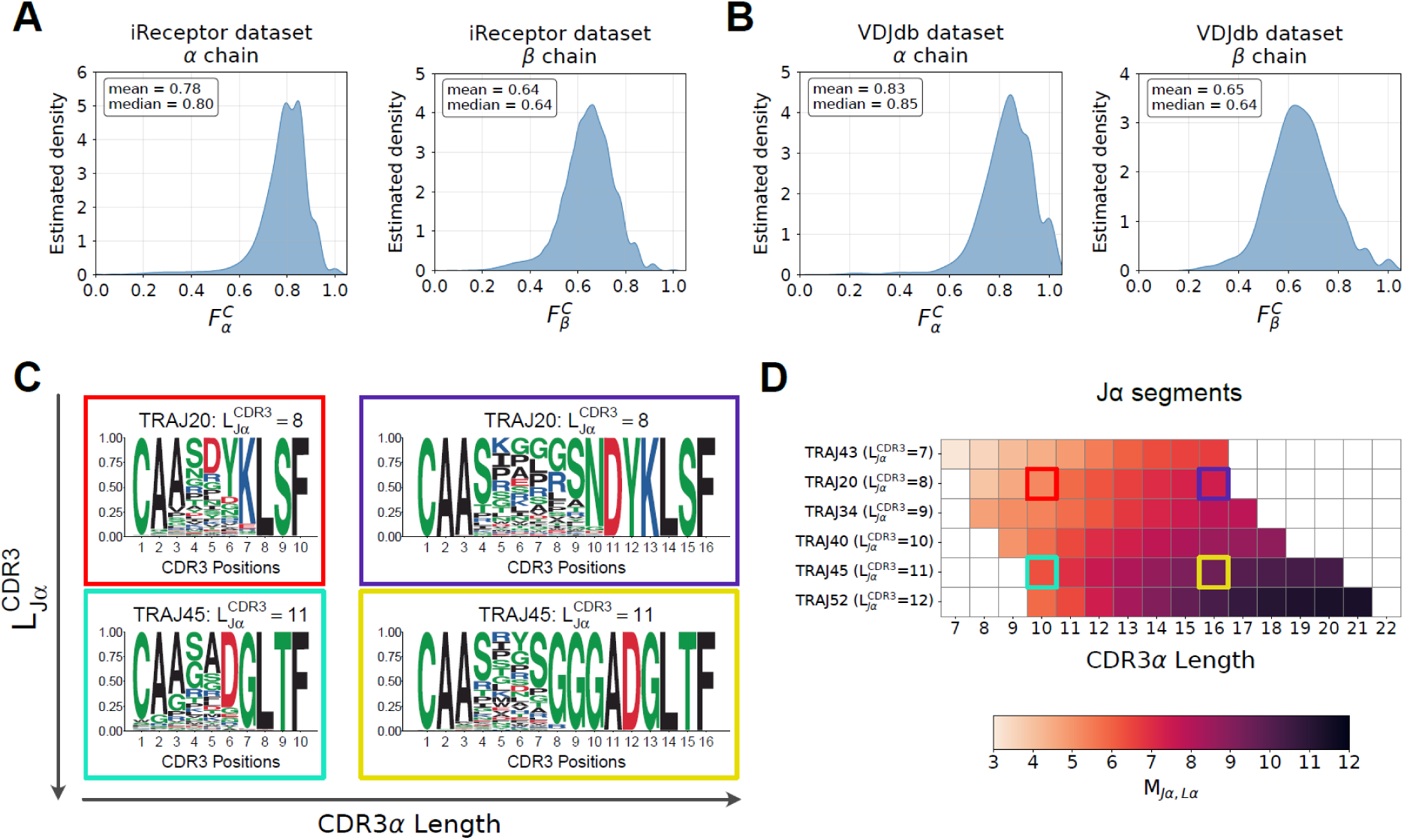
**V and J gene usage shapes the amino acid composition of CDR3 sequences**. **(A)** Fraction (F^C^_α_ and F^C^_β_) of the CDR3 residues that can be mapped to germline sequences of V and J genes in TCR repertoires of unknown specificity (iReceptor dataset). **(B)** Fraction (F^C^_α_ and F^C^_β_) of CDR3 residues that can be mapped to germline sequences of V and J genes in epitope-specific TCR repertoires (VDJdb dataset). **(C)** Examples of CDR3α motifs of different lengths (10 or 16) derived from TCRs with a short Jα gene (TRAJ20, L^CDR3^_TRAJ20_ = 8, `SNDYKLSF`) or a long Jα gene (TRAJ45, L^CDR3^_TRAJ45_ = 11, `YSGGGADGLTF`), and the same Vα gene (TRAV13-1). **(D)** Mean number of conserved germline amino acids (M_Jα,Lα_) for selected Jα genes with distinct L^CDR3^_jβ_ values (see Supp. Figure 3 for all M values in human and Supp. Figure 4 for all M_V/J,L_ values in murine). Colored squares refer to the examples in panel C.

Overall, our results demonstrate that the number of CDR3 residues in germline V and J sequences has a profound impact on CDR3 lengths in TCR repertoires of undetermined or known specificity.

### V and J gene usage shapes the amino acid composition of CDR3 sequences

We next explored the impact of V and J genes on the CDR3 amino acid composition. First, for each TCR, we calculated the number of CDR3 residues that could be mapped to the germline sequences of the V and J genes. To this end, we counted the number of consecutive residues at the beginning and end of each CDR3 that matched their respective germline sequences and divided by the length of the corresponding CDR3. We denoted this quantity by F^C^_α_ and F^C^_β_. The estimated density of the F^C^_α_ and F^C^_β_ values in our collection of TCR repertoires with undetermined specificity showed that on average 80% of CDR3α and 65% of CDR3β residues could be mapped to germline V and J sequences (Figure 2A). Similar results were obtained in TCR repertoires with known specificity (Figure 2B).

Next, we investigated how each combination of germline V and J genes shapes the amino acid composition of CDR3 sequences across different CDR3 lengths. To illustrate this analysis, we first examined motifs of CDR3α of length 10 and 16 derived from TCRs with the same Vα gene (TRAV13-1) and either a short Jα gene (TRAJ20, `SNDYKLSF`, L^CDR3^_TRAJ20_ = 8) or a long Jα gene (TRAJ45, `YSGGGADGLTF`, L^CDR3^_TRAJ45_ = 11) (Figure 2C). For the short Jα gene (TRAJ20), CDR3s of length 10 retained the final three germline residues (`LSF`), whereas CDR3s of length 16 retained the final six germline residues (`DYKLSF`). For the longer Jα gene (TRAJ45), CDR3s of length 10 retained the final four germline residues (`GLTF`), while those of length 16 retained the final eight germline residues (`GGADGLTF`).

Overall, these examples suggest that the average number of CDR3 residues determined by V and J usage depends both on the CDR3 length and the length of the V and J genes (i.e., L^CDR3^_v/j_), with higher values for longer CDR3s and higher L^CDR3^_v/j_. To systematically characterize this effect, we computed for each Vα, Jα, Vβ and Jβ, and for each CDR3 length (L) the mean number of consecutive residues matching the germline sequence, denoted as M_V/J,L_ (Supp. Figure 3, see Supp. Figure 4 for murine data and Figure 2D for selected examples with Jα genes representative of the different values of L^CDR3^_vα_ in Human). M increased with both L^CDR3^_v/j_ and CDR3 lengths, with the most pronounced effect observed for Jα genes.

**Figure 3.**
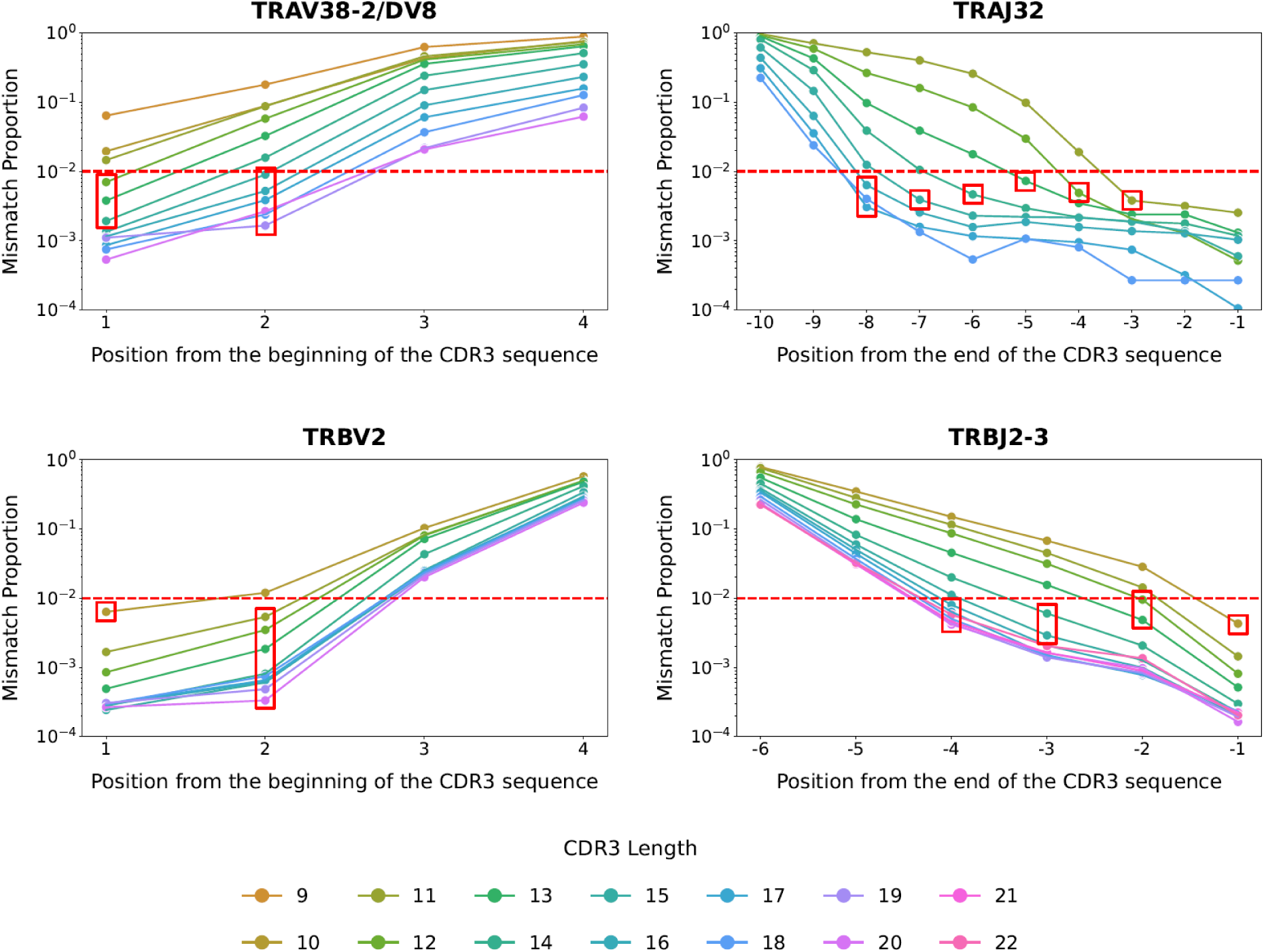
Proportion of mismatches at different CDR3 positions for selected V and J genes. Each dot represents the proportion of CDR3 sequences with a non-germline amino acid at the corresponding position. The red dashed line indicates the 0.01 mismatch threshold used to define the expected number of consistently conserved germline residues (i.e., T_V/J,L_). The red boxes indicate, for each line, the last position below the threshold of 0.01.

**Figure 4:**
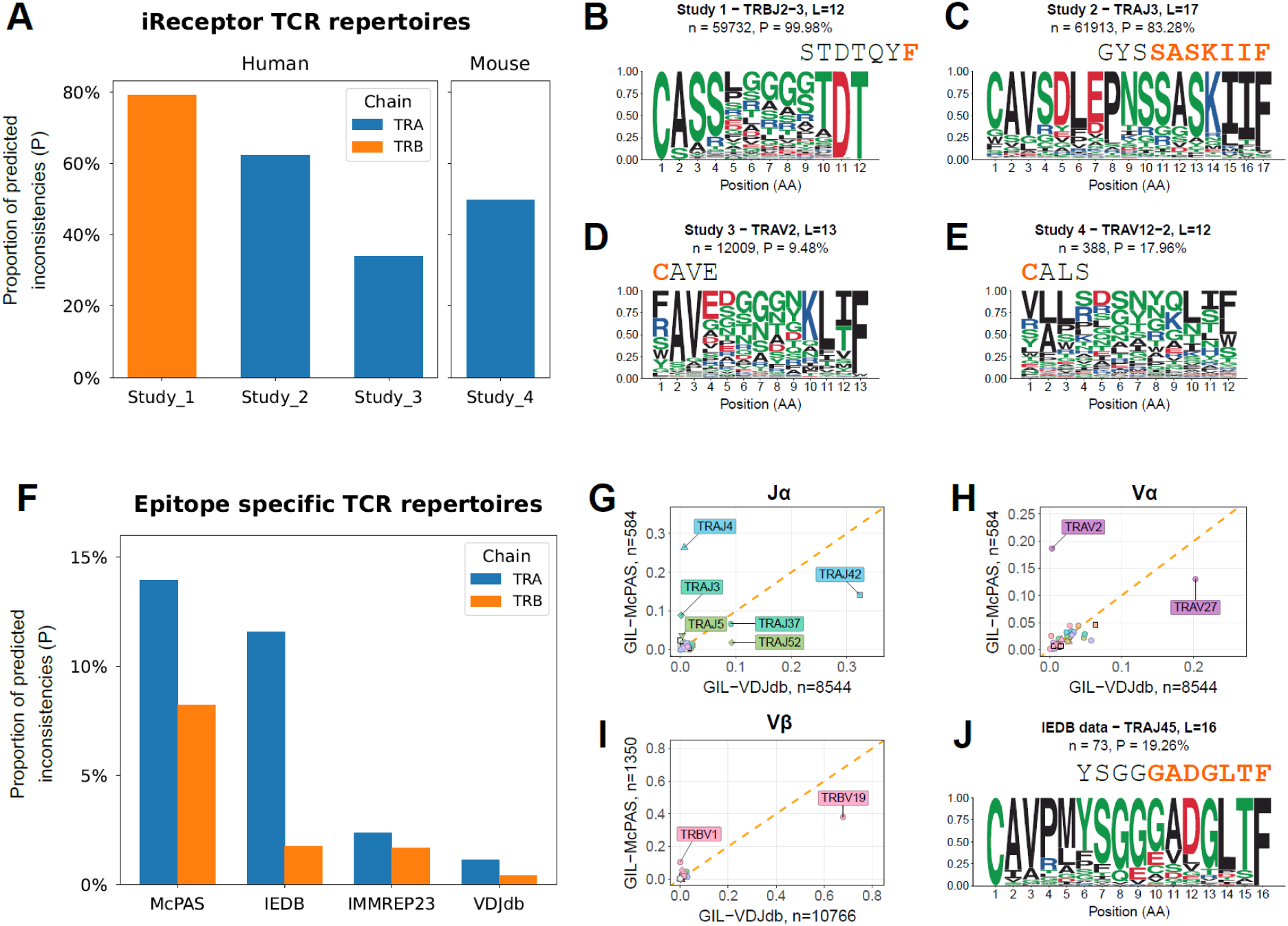
Consistency between V and J gene usage and CDR3 sequences provides a robust quality control for TCR repertoire datasets. **(A)** Proportion (P) of sequences predicted as inconsistent in TCR repertoires of unknown specificity from four studies in the iReceptor database. **(B)** Example of systematic truncation of C-terminal residues in CDR3β of TRBJ2-3 TCRs predicted as inconsistent (Study 1) (‘n’ stands for the number of predicted inconsistencies, and ‘P’ for their proportion among all TRBJ2-3 TCRs). **(C)** Example of widespread predicted inconsistencies across CDR3α positions in TRAJ3 TCRs (Study 2). **(D)** Example of loss of the N-terminal cysteine in CDR3α sequences of TRAV2 TCRs predicted as inconsistent (Study 3). **(E)** Example of loss of the N-terminal cysteine in CDR3α sequences of murine TRAV12-2 TCRs predicted as inconsistent (Study 4). **(F)** Proportion of predicted inconsistent sequences across four TCR repertoire datasets with known specificity (McPAS, IEDB, IMMREP23, and VDJdb). **(G)** Comparison of Jα gene usage frequencies between McPAS and VDJdb for GIL-specific TCRs. **(H)** Comparison of Vα gene usage frequencies between McPAS and VDJdb for GIL-specific TCRs. **(I)** Comparison of Vβ gene usage frequencies between McPAS and VDJdb for GIL-specific TCRs. **(J)** Motifs of CDR3α sequences of TRAJ45 TCRs in IEDB predicted to be inconsistent. Orange letters in panels B, C, D, E and J represent the germline amino acid expected to be conserved based on the T_V,L_ and T_J,L_ thresholds.

Altogether, our findings demonstrate that on average 80% of the CDR3α and 65% of the CDR3β residues are determined by the choices of V and J genes, and that longer CDR3 sequences and V and J genes with a higher number of CDR3 residues correlate with higher numbers of CDR3 residues matching those of the germline V and J genes.

### Consistency between V and J gene usage and CDR3 sequences provides a robust quality control for TCR repertoire datasets

We next hypothesized that our results could provide a useful quality control approach for TCR repertoire datasets by investigating consistency (or lack thereof) between V and J gene annotations and CDR3 sequences. To this end, we first calculated how often the amino acid at each CDR3 position differs from the one encoded by the corresponding germline V or J gene in our collection of TCR repertoires. These computations were performed separately for each V and J gene and each CDR3 length. As expected, mismatch frequencies were lowest at the beginning and the end of the CDR3s and gradually increased toward the middle of these sequences (Figure 3). Consistent with the previous analyses, longer CDR3s generally exhibited higher conservation at comparable positions. We then determined the number of residues that were consistently conserved by counting the number of consecutive positions whose mismatch proportion remained below 0.01. This procedure was performed for each CDR3 length and each Vα, Jα, Vβ and Jβ and the resulting numbers of conserved residues were denoted as T_Vα,Lα_, T_Jα,Lα_, T_Vβ,Lβ_ and T_Jβ,Lβ_ (see Methods, Supp. Figure 5 and red boxes in the examples of Figure 3). These thresholds values were used to investigate potential issues in TCR repertoire datasets by evaluating for each TCR sequence whether the expected number of germline-encoded residues (i.e., T_V,L_ and T_J,L_) was found at the beginning and end of CDR3s. TCR sequences that deviate from this criterion are referred to as “predicted inconsistencies”.

To illustrate the capability of this approach to detect important issues in TCR repertoire datasets, we first applied it to multiple human and murine TCR repertoires of unknown specificity and selected four datasets with high proportions of predicted inconsistencies (Figure 4A, Supp. Table 3).

In the first example (Study 1, human TCRβ data), nearly all sequences were classified as inconsistent. Examination of the predicted inconsistent sequences revealed a systematic truncation, with the last two or the last three CDR3 amino acids missing (see Figure 4B for TRBJ2-3 and Supp. Figure 6A for TRBJ1-1 and TRBJ2-1). The second example (Study 2, human TCRα data) showed an unexpected level of predicted inconsistencies at many CDR3 positions. For instance, CDR3 sequences of length 17 from TRAJ3 TCRs with predicted inconsistencies displayed the expected germline motif (`SSASKIIF`), but all positions contained high proportions of non-germline amino acids (Figure 4C). Finally, Studies 3 and 4 highlight cases where predicted inconsistencies lacked the canonical N-terminal cysteine in human (Figure 4D) and murine (Figure 4E) TCRs. For these reasons, none of these studies was included in the collection of TCR repertoires used throughout this manuscript.

To investigate whether similar issues could arise in datasets of epitope-specific TCRs, we applied our approach to four epitope-specific datasets, including McPAS (Tickotsky et al., 2017), IEDB (Vita et al., 2015), IMMREP23 (Nielsen et al., 2024), and VDJdb (Shugay et al., 2018) and quantified the proportion of predicted inconsistencies in TCRα and TCRβ sequences (Figure 4F). The McPAS database exhibited the highest fraction of predicted inconsistencies (14% in TCRα, 8% in TCRβ). To explore potential issues in this database, we focused on TCRs specific for the well-studied influenza epitope `GILGFVFTL` restricted to HLA-A*02:01 (GIL), which displayed one of the highest proportions of predicted inconsistencies. Comparison of GIL-specific repertoires between McPAS and VDJdb revealed major discrepancies in V and J gene usage. In VDJdb, GIL-specific TCRα sequences were dominated by TRAJ42 (see also https://tcrmotifatlas.unil.ch/model/A0201_GILGFVFTL and (Liu et al., 2025)), whereas in McPAS TRAJ4 was the most frequent Jα gene (Figure 4G). Additional discrepancies were observed between TRAJ37 and TRAJ3 and between TRAJ52 and TRAJ5. For each GIL-specific TRAJ4 TCR predicted as inconsistent, we determined the Jα gene best compatible with the CDR3α. This analysis revealed that all CDR3 sequences were compatible with some TRAJ4x genes (x=0,…,9, Supp. Figure 6B, with >70% TRAJ42), suggesting that the last digit of the Jα gene names may have been lost. We next investigated whether this issue could also affect other V or J genes. We observed that TRAV2 was the most frequent V gene in GIL-specific TCRs in the McPAS dataset, whereas TRAV27 was the most frequent in VDJdb (Figure 4H). Likewise, the pseudogene TRBV1 was considerably overrepresented in McPAS, while TRBV19 was underrepresented relative to VDJdb (Figure 4I). Inspection of the CDR3 sequences for TCRs annotated as TRAV2 and TRBV1 in McPas confirmed better consistency with TRAV27 and TRBV19, respectively (Supp. Figure 6C-E). Collectively, these findings suggest that a fraction of TCRs in the McPAS database may suffer from a systematic post-processing error in which the second digit of the V and J gene names was inadvertently removed.

In the case of the IEDB dataset, a high proportion of TCRα sequences were classified as inconsistent. To illustrate this observation, the CDR3 motif of TRAJ45 TCR sequences showed a high frequency of amino acids different from those of TRAJ45 at positions expected to show very few non-germline residues (Figure 4J). Regarding the IMMREP23 datasets, our approach recapitulated the issue discussed during the competition (i.e., missing F in some CDR3β of Jβ genes with two consecutive F residues at the end of the CDR3).

Overall, these examples illustrate how our quantification of the impact of V and J gene usage on CDR3 sequences can pinpoint specific issues in TCR repertoire datasets, including some that have been widely used to benchmark TCR-epitope prediction algorithms.

## Discussion

TCRs are characterized by a high sequence diversity that reflects both the different choices of V and J genes and the rearrangements taking place at the V(D)J junctions within the CDR3 loops of both chains. For these reasons, TCR sequences are often represented by the V and J gene names and the sequence of the CDR3s. However, the extent to which V and J gene usage influences CDR3 sequences has not been systematically investigated across all V and J genes and CDR3 lengths.

In this work, we precisely determined how the length and the amino acid composition of CDR3s are impacted by the V and J choices. Our results show that a key determinant of CDR3 lengths can be traced back to the number of CDR3 residues that exist in germline V and J genes. Our work further indicates that on average 80% of CDR3α residues and 65% of CDR3β residues are determined by V and J gene usage in TCR repertoires of either undetermined or known specificity. In our analyses, we did not consider the two D genes (TRBD1-`GTGG` and TRBD2-`GTSGG`) in TCRβ chains, since D gene usage is not reported in many datasets and can be challenging to unambiguously determine from TCR sequencing data (Omer et al., 2022).

We anticipate that the presence of residues encoded by the D gene in CDR3β will further increase the fraction of CDR3β residues determined by germline-encoded genes.

These findings challenge a view sometimes held that variations in CDR3 length and amino acid composition would primarily reflect nucleotide insertions and deletions at the V(D)J junctions. In particular, our results indicate that CDR3 sequences should not be treated as protein sequences freely acquiring mutations at different positions independently of the V and J genes. For instance, any constraint (e.g., recognition of a specific epitope (Dash et al., 2017; Liu et al., 2025)) imposing specific amino acids in CDR1/2 loops will impact V gene usage, which will in turn create specific amino acid patterns at the beginning of CDR3 sequences. Similarly, requirements for a specific amino acid at a given position towards the end of CDR3 sequences will impose constraints on J gene usage, which will in turn impact many other CDR3 residues as well as CDR3 lengths. This also means that batch effects or biases resulting in different V and J gene usage across different TCR repertoires (Barennes et al., 2021) will influence statistical properties of CDR3 sequences and can result in apparent enrichment of specific CDR3 motifs or k-mers. Overall, our results emphasize how disregarding the profound impact of V and J gene usage on CDR3 sequences can lead to multiple confounding factors when interpreting CDR3 sequence patterns in different contexts.

Our work further demonstrates how a precise quantification of the number of CDR3 residues unlikely to differ from those expected in the germline V and J genes can help highlight potential issues in some TCR repertoire datasets. These issues comprise putative problems in TCR sequencing, TCR reconstruction (including errors in V/J annotation or variable definitions of CDR3s within the same study) or data processing. To predict such inconsistencies, we used a threshold of 1% on the false-discovery rate. For this reason, we cannot assume that any TCR sequence with a CDR3 that does not match the expectations from V and J gene choices is necessarily problematic. In addition, we cannot exclude that some inconsistencies could originate from rare alleles of V or J genes with distinct CDR3 residues that are not included in the IMGT reference used in this work. However, the different examples described in this work show that our approach is useful for detecting different sources of potential issues in TCR repertoire datasets that affect a high (i.e., much higher than 1%) fraction of TCRs. To infer the thresholds T_V,L_ and T_J,L_ we used TCR repertoires which could themselves contain different sources of noise. This could result in slightly underestimating these thresholds. Despite this limitation, our proposed approach provides a way to pinpoint several issues in widely used TCR repertoire datasets that could not have been detected by existing tools (Nagano and Chain, 2023).

Altogether, our results demonstrate that the germline sequences of the V and J genes play a major role in shaping CDR3 length and amino acid composition. Building on these findings, we developed a framework to identify putative inconsistencies between annotated V and J genes and CDR3 sequences. This method available through the MixTCRclean R package (https://github.com/GfellerLab/MixTCRclean). Applying this approach to different TCR repertoires reveals important issues in several datasets and supports the idea that consistency analysis between CDR3 sequences and V and J gene usage can be used as a quality control for TCR repertoires.

## Methods

### Collection and curation of TCR repertoires

#### Human data

We collected bulk T-cell receptor (TCR) repertoire datasets (Supp. Table 1) for both the α and β chains from the iReceptor platform (Corrie et al., 2018). To ensure consistency and data quality, we applied a series of preprocessing steps. CDR3 sequences containing non-standard amino acids were removed, and only those between 7 and 22 amino acids in length were retained. Allele-level information was discarded when provided, and gene names were standardized according to IMGT nomenclature. Only TCRs with complete V, J, and CDR3 information were kept, and a single clone per subject repertoire was retained. Finally, subjects with fewer than 1,000 TCR sequences were excluded.

#### Murine data

Murine bulk TCR repertoires were collected from three sources: the Adaptive Biotech website and two other studies (Brown et al., 2024; Genolet et al., 2023) (see Supp. Table 2 for details).

Preprocessing steps were largely consistent with those applied to human datasets, with two exceptions. First, due to the smaller number of available TCRs, all murine repertoires were retained, including those with low sequence counts. Second, V and J gene names corresponding to different murine strains were merged to ensure consistent annotation across datasets.

### Epitope-specific data

Epitope-specific TCR sequences were obtained from four publicly available databases, all downloaded on August 4, 2025: IEDB (human TCR sequences only) (Vita et al., 2019), VDJdb (human TCR sequences only) (Shugay et al., 2018), McPAS-TCR (human TCR sequences only) (Tickotsky et al., 2017), and IMMREP23 (Nielsen et al., 2024), using the *solution.csv* file from the IMMREP23 GitHub repository (https://github.com/justin-barton/IMMREP23).

For VDJdb, McPAS, and IMMREP23, minimal preprocessing was applied following the same procedures as for the iReceptor human data described above.

Regarding IEDB data, additional curation was performed to harmonize CDR3 boundary definitions with IMGT standards. A subset of IEDB sequences appeared to use an alternative definition where the CDR3 region begins after the conserved cysteine and ends one amino acid before the standard IMGT endpoint. For these sequences, we restored the missing germline residues by adding the corresponding first and last (or last two) amino acids from the annotated V and J genes.

### Fraction of conserved germline residues: F^C^ and F^C^

For each TCR in the human iReceptor dataset and the human VDJdb dataset, we computed the fraction of conserved germline residues (F^C^_α_ and F^C^_β_). For a given sequence, we first counted the number of consecutive residues from the beginning of the CDR3 that matched the germline amino acid sequence of the annotated V gene. We then repeated this from the end of the CDR3 against the germline sequence of the annotated J gene. The two counts were summed and divided by the CDR3 length to obtain F^C^_α/β_. When multiple alleles existed for an annotated V or J gene, we matched the CDR3 against each allele’s germline sequence and retained the maximum number of consecutive matches. Kernel density estimates in Figures 2A and 2B were generated using the Seaborn library (`kdeplot`) with a Gaussian kernel.

### Mean number of conserved germline amino acids: M_V,L_ and M_J,L_

As described above, we counted the number of consecutive germline-encoded amino acids at the beginning (V gene) and end (J gene) of each CDR3 that matched the corresponding germline reference, treating multiple alleles as before (i.e., using the allele that yielded the maximum consecutive match). For each combination of genes (V or J) and CDR3 length, we then computed the mean of these counts across all sequences, yielding M_Vα,Lα_, M_Jα,Lα,_ M_Vβ,Lβ_ and M_Jβ,Lβ_. To ensure robust estimates, we included only combinations represented by at least 1,000 TCRα sequences or 10,000 TCRβ sequences. Consequently, very rare genes and sequences with extremely short or long CDR3s were excluded. For the murine datasets, where fewer sequences were available, the same statistics were computed for combinations with at least 500 sequences (α or β).

### Inference of position-wise mismatch proportion and computation of T_V,L_ and T_J,L_

We applied the same minimum data thresholds used for M_V/J,L_. For each combination of V and J genes and CDR3 length, and for each CDR3 position that could be influenced by the germline V or J sequence, we identified sequences carrying a mismatch relative to the corresponding germline residue. To distinguish true mismatches from potential simple sequencing errors, each amino-acid mismatch was examined at the nucleotide level. A potential simple sequencing error was defined as a single-nucleotide substitution isolated within a ±3 nucleotide window. Concretely, for a mismatch at nucleotide position i, we required exact matches at positions i-3, i-2, i-1 and i+1, i+2, i+3. If this criterion was met, the event was classified as a potential sequencing error and excluded. We treated genes with multiple alleles as before (i.e. allowing a match to any allele’s germline sequence).

The position-wise mismatch proportion for each gene (V or J) and each CDR3 length was then defined as the fraction of sequences with a germline mismatch not attributable to simple sequencing errors.

We then set a threshold of 0.01 and counted for each gene (V or J)and CDR3 length the number of consecutive positions for which the position-wise mismatch proportions were lower than this threshold from the beginning and end of the CDR3 sequences. This gave us the number of residues that are consistently conserved from the germline sequences of the V and J genes with a false discovery rate of 1%. These initial estimates were limited to V and J genes and CDR3 length combinations with sufficient sequence counts. To extend this approach to any TCR sequence, we generalized these quantities by leveraging the strong dependence of germline conservation on L^CDR3^_v/j_. We grouped all V and J genes within each family by their nucleotide length and computed the mean of the available T_V,L_ and T_J,L_ values for each group. The resulting points were used to perform a piecewise linear interpolation describing the relationship between CDR3 length and the expected number of conserved residues. To yield integer counts of conserved residues, the interpolated results were rounded down to the nearest integer for each CDR3 length.

### Prediction of inconsistencies between V and J genes and CDR3 sequences

To predict inconsistencies, we used the thresholds T_V,L_ and T_J,L_. For the beginning of each CDR3, we compared the first T_V,L_ residues to the annotated V germline sequence (considering all alleles as described above). For the end, we compared the last T_J,L_ residues to the annotated J germline sequence (again considering all alleles). If any mismatch was detected within either required substring, the sequence was annotated as inconsistent; otherwise, it was annotated as consistent.

We integrated this framework into an R package called MixTCRclean which provides standardization and pre-cleaning steps as well as prediction of inconsistencies.

## Data availability

All datasets used in this study were collected from publicly available repositories. Detailed sources for human and murine TCR repertoires of unknown specificity are listed in Supplementary Tables 1-3. Epitope-specific TCR repertoires were downloadedon August 4, 2025 from VDJdb (https://vdjdb.cdr3.net/), IEDB (https://www.iedb.org/), McPAS

(https://friedmanlab.weizmann.ac.il/McPAS-TCR/) and IMMREP23 (https://github.com/justin-barton/IMMREP23).

## Code availability

The MixTCRclean package is freely available at https://github.com/GfellerLab/MixTCRclean.

## References

1. Alamyar, E., Giudicelli, V., Li, S., Duroux, P., Lefranc, M.-P., 2012. IMGT/HighV-QUEST: the IMGT® web portal for immunoglobulin (IG) or antibody and T cell receptor (TR) analysis from NGS high throughput and deep sequencing. Immunome Res. 8, 26. 10.4172/1745-7580.1000056

2. Bai, H., Ma, J., Mao, W., Zhang, X., Nie, Y., Hao, J., Wang, X., Qin, H., Zeng, Q., Hu, F., Qi, X., Chen, X., Li, D., Zhang, B., Shi, B., Zhang, C., 2022. Identification of TCR repertoires in asymptomatic COVID-19 patients by single-cell T-cell receptor sequencing. Blood Cells. Mol. Dis. 97, 102678. 10.1016/j.bcmd.2022.102678

3. Barennes, P., Quiniou, V., Shugay, M., Egorov, E.S., Davydov, A.N., Chudakov, D.M., Uddin, I., Ismail, M., Oakes, T., Chain, B., Eugster, A., Kashofer, K., Rainer, P.P., Darko, S., Ransier, A., Douek, D.C., Klatzmann, D., Mariotti-Ferrandiz, E., 2021. Benchmarking of T cell receptor repertoire profiling methods reveals large systematic biases. Nat. Biotechnol. 39, 236–245. 10.1038/s41587-020-0656-3

4. Beshnova, D., Ye, J., Onabolu, O., Moon, B., Zheng, W., Fu, Y.-X., Brugarolas, J., Lea, J., Li, B., 2020. De novo prediction of cancer-associated T cell receptors for noninvasive cancer detection. Sci. Transl. Med. 12, eaaz3738. 10.1126/scitranslmed.aaz3738

5. Bolotin, D.A., Mamedov, I.Z., Britanova, O.V., Zvyagin, I.V., Shagin, D., Ustyugova, S.V., Turchaninova, M.A., Lukyanov, S., Lebedev, Y.B., Chudakov, D.M., 2012. Next generation sequencing for TCR repertoire profiling: platform-specific features and correction algorithms. Eur. J. Immunol. 42, 3073–3083. 10.1002/eji.201242517

6. Bolotin, D.A., Poslavsky, S., Mitrophanov, I., Shugay, M., Mamedov, I.Z., Putintseva, E.V., Chudakov, D.M., 2015. MiXCR: software for comprehensive adaptive immunity profiling. Nat. Methods 12, 380–381. 10.1038/nmeth.3364

7. Bradley, P., Thomas, P.G., 2019. Using T Cell Receptor Repertoires to Understand the Principles of Adaptive Immune Recognition. Annu. Rev. Immunol. 37, 547–570. 10.1146/annurev-immunol-042718-041757

8. Brown, A.J., White, J., Shaw, L., Gross, J., Slabodkin, A., Kushner, E., Greiff, V., Matsuda, J., Gapin, L., Scott-Browne, J., Kappler, J., Marrack, P., 2024. MHC heterozygosity limits T cell receptor variability in CD4 T cells. Sci. Immunol. 9, eado5295. 10.1126/sciimmunol.ado5295

9. Charles, J., Mouret, S., Challende, I., Leccia, M.-T., De Fraipont, F., Perez, S., Plantier, N., Plumas, J., Manuel, M., Chaperot, L., Aspord, C., 2020. T-cell receptor diversity as a prognostic biomarker in melanoma patients. Pigment Cell Melanoma Res. 33, 612–624. 10.1111/pcmr.12866

10. Corrie, B.D., Marthandan, N., Zimonja, B., Jaglale, J., Zhou, Y., Barr, E., Knoetze, N., Breden, F.M.W., Christley, S., Scott, J.K., Cowell, L.G., Breden, F., 2018. iReceptor: A platform for querying and analyzing antibody/B-cell and T-cell receptor repertoire data across federated repositories. Immunol. Rev. 284, 24–41. 10.1111/imr.12666

11. Dash, P., Fiore-Gartland, A.J., Hertz, T., Wang, G.C., Sharma, S., Souquette, A., Crawford, J.C., Clemens, E.B., Nguyen, T.H.O., Kedzierska, K., La Gruta, N.L., Bradley, P., Thomas, P.G., 2017. Quantifiable predictive features define epitope-specific T cell receptor repertoires. Nature 547, 89–93. 10.1038/nature22383

12. DeWitt, W.S., Emerson, R.O., Lindau, P., Vignali, M., Snyder, T.M., Desmarais, C., Sanders, C., Utsugi, H., Warren, E.H., McElrath, J., Makar, K.W., Wald, A., Robins, H.S., 2015. Dynamics of the cytotoxic T cell response to a model of acute viral infection. J. Virol. 89, 4517–4526. 10.1128/JVI.03474-14

13. Dong, N., Moreno-Manuel, A., Calabuig-Fariñas, S., Gallach, S., Zhang, F., Blasco, A., Aparisi, F., Meri-Abad, M., Guijarro, R., Sirera, R., Camps, C., Jantus-Lewintre, E., 2021. Characterization of Circulating T Cell Receptor Repertoire Provides Information about Clinical Outcome after PD-1 Blockade in Advanced Non-Small Cell Lung Cancer Patients. Cancers 13, 2950. 10.3390/cancers13122950

14. Dudgeon, C., Chan, C., Kang, W., Sun, Y., Emerson, R., Robins, H., Levine, A.J., 2014. The evolution of thymic lymphomas in p53 knockout mice. Genes Dev. 28, 2613–2620. 10.1101/gad.252148.114

15. Dupic, T., Marcou, Q., Walczak, A.M., Mora, T., 2019. Genesis of the αβ T-cell receptor. PLOS Comput. Biol. 15, e1006874. 10.1371/journal.pcbi.1006874

16. Freeman, J.D., Warren, R.L., Webb, J.R., Nelson, B.H., Holt, R.A., 2009. Profiling the T-cell receptor beta-chain repertoire by massively parallel sequencing. Genome Res. 19, 1817–1824. 10.1101/gr.092924.109

17. Gaide, O., Emerson, R.O., Jiang, X., Gulati, N., Nizza, S., Desmarais, C., Robins, H., Krueger, J.G., Clark, R.A., Kupper, T.S., 2015. Common clonal origin of central and resident memory T cells following skin immunization. Nat. Med. 21, 647–653. 10.1038/nm.3860

18. Genolet, R., Bobisse, S., Chiffelle, J., Arnaud, M., Petremand, R., Queiroz, L., Michel, A., Reichenbach, P., Cesbron, J., Auger, A., Baumgaertner, P., Guillaume, P., Schmidt, J., Irving, M., Kandalaft, L.E., Speiser, D.E., Coukos, G., Harari, A., 2023. TCR sequencing and cloning methods for repertoire analysis and isolation of tumor-reactive TCRs. Cell Rep. Methods 3, 100459. 10.1016/j.crmeth.2023.100459

19. Giudicelli, V., 2004. IMGT/GENE-DB: a comprehensive database for human and mouse immunoglobulin and T cell receptor genes. Nucleic Acids Res. 33, D256–D261. 10.1093/nar/gki010

20. Greissl, J., Pesesky, M., Dalai, S.C., Rebman, A.W., Soloski, M.J., Horn, E.J., Dines, J.N., Gill, D.B., Gittelman, R.M., Snyder, T.M., Emerson, R.O., Meeds, E., Manley, T., Kaplan, I.M., Baldo, L., Carlson, J.M., Robins, H.S., Aucott, J.N., 2022. Immunosequencing of the T-Cell Receptor Repertoire Reveals Signatures Specific for Identification and Characterization of Early Lyme Disease. 10.1101/2021.07.30.21261353

21. Hannan, R., Mohamad, O., Diaz de Leon, A., Manna, S., Pop, L.M., Zhang, Z., Mannala, S., Christie, A., Christley, S., Monson, N., Ishihara, D., Hsu, E.J., Ahn, C., Kapur, P., Chen, M., Arriaga, Y., Courtney, K., Cantarel, B., Wakeland, E.K., Fu, Y.-X., Pedrosa, I., Cowell, L., Wang, T., Margulis, V., Choy, H., Timmerman, R.D., Brugarolas, J., 2021. Outcome and Immune Correlates of a Phase II Trial of High-Dose Interleukin-2 and Stereotactic Ablative Radiotherapy for Metastatic Renal Cell Carcinoma. Clin. Cancer Res. Off. J. Am. Assoc. Cancer Res. 27, 6716–6725. 10.1158/1078-0432.CCR-21-2083

22. Heather, J.M., Best, K., Oakes, T., Gray, E.R., Roe, J.K., Thomas, N., Friedman, N., Noursadeghi, M., Chain, B., 2016. Dynamic Perturbations of the T-Cell Receptor Repertoire in Chronic HIV Infection and following Antiretroviral Therapy. Front. Immunol. 6, 644. 10.3389/fimmu.2015.00644

23. Heather, J.M., Ismail, M., Oakes, T., Chain, B., 2017. High-throughput sequencing of the T-cell receptor repertoire: pitfalls and opportunities. Brief. Bioinform. 19, 554–565. 10.1093/bib/bbw138

24. Iijima, N., Iwasaki, A., 2014. A local macrophage chemokine network sustains protective tissue-resident memory CD4 T cells. Science 346, 93–98. 10.1126/science.1257530

25. Jia, Q., Wu, W., Wang, Y., Alexander, P.B., Sun, C., Gong, Z., Cheng, J.-N., Sun, H., Guan, Y., Xia, X., Yang, L., Yi, X., Wan, Y.Y., Wang, H., He, J., Futreal, P.A., Li, Q.-J., Zhu, B., 2018. Local mutational diversity drives intratumoral immune heterogeneity in non-small cell lung cancer. Nat. Commun. 9, 5361. 10.1038/s41467-018-07767-w

26. Jiang, Y., Huo, M., Cheng Li, S., 2023. TEINet: a deep learning framework for prediction of TCR-epitope binding specificity. Brief. Bioinform. 24, bbad086. 10.1093/bib/bbad086

27. Joseph, M., Wu, Y., Dannebaum, R., Rubelt, F., Zlatareva, I., Lorenc, A., Du, Z.G., Davies, D., Kyle-Cezar, F., Das, A., Gee, S., Seow, J., Graham, C., Telman, D., Bermejo, C., Lin, H., Asgharian, H., Laing, A.G., del Molino del Barrio, I., Monin, L., Muñoz-Ruiz, M., McKenzie, D.R., Hayday, T.S., Francos-Quijorna, I., Kamdar, S., Davis, R., Sofra, V., Cano, F., Theodoridis, E., Martinez, L., Merrick, B., Bisnauthsing, K., Brooks, K., Edgeworth, J., Cason, J., Mant, C., Doores, K.J., Vantourout, P., Luong, K., Berka, J., Hayday, A.C., 2022. Global patterns of antigen receptor repertoire disruption across adaptive immune compartments in COVID-19. Proc. Natl. Acad. Sci. 119, e2201541119. 10.1073/pnas.2201541119

28. Lang Kuhs, K.A., Lin, S.-W., Hua, X., Schiffman, M., Burk, R.D., Rodriguez, A.C., Herrero, R., Abnet, C.C., Freedman, N.D., Pinto, L.A., Hamm, D., Robins, H., Hildesheim, A., Shi, J., Safaeian, M., 2018. T cell receptor repertoire among women who cleared and failed to clear cervical human papillomavirus infection: An exploratory proof-of-principle study. PLoS ONE 13, e0178167. 10.1371/journal.pone.0178167

29. Levi, R., Louzoun, Y., 2022. Two Step Selection for Bias in β Chain V-J Pairing. Front. Immunol. 13.

30. Li, Y., Nahas, M., Stephens, D., Froburg, K., Hintz, E., Champagne, D., Lochab, A., Brown, M., Braun, J., Fortuño, M.A., Ocón, M.-M., Pasquier, A., Luque-Vázquez, I., Moudgalya, H., Kivlehan, S., Gjeci, I., Korle, S.L., Campo, A., Rodriguez, M., Seder, C.W., Lizotte, P.H., Bueno, R., Borgia, J.A., Seijo, L.M., Montuenga, L.M., Yelensky, R., 2025. Circulating T-cell receptor repertoire for cancer early detection. Npj Precis. Oncol. 9, 245. 10.1038/s41698-025-01036-y

31. Liu, P., Liu, D., Yang, X., Gao, J., Chen, Y., Xiao, X., Liu, F., Zou, J., Wu, J., Ma, J., Zhao, F., Zhou, X., Gao, G.F., Zhu, B., 2014. Characterization of human αβTCR repertoire and discovery of D-D fusion in TCRβ chains. Protein Cell 5, 603–615. 10.1007/s13238-014-0060-1

32. Liu, Y., Croce, G., Tadros, D., Moreno, D., Michel, A., Thierry, A.-C., Genolet, R., Perez, M.A., Lani, R., Guillaume, P., Hebeisen, M., Teleman, M., Lau, K., Larabi, A., Racle, J., Speiser, D.E., Pojer, F., Dunn, S., Baumgaertner, P., Zoete, V., Harari, A., Gfeller, D., 2025. Key determinants of T cell epitope recognition revealed by TCR specificity profiles. 10.1101/2025.11.17.688817

33. Ma, L., Yang, L., Bin Shi, He, X., Peng, A., Li, Y., Zhang, T., Sun, S., Ma, R., Yao, X., 2016. Analyzing the CDR3 Repertoire with respect to TCR—Beta Chain V-D-J and V-J Rearrangements in Peripheral T Cells using HTS. Sci. Rep. 6, 29544. 10.1038/srep29544

34. Maceiras, A.R., Almeida, S.C.P., Mariotti-Ferrandiz, E., Chaara, W., Jebbawi, F., Six, A., Hori, S., Klatzmann, D., Faro, J., Graca, L., 2017. T follicular helper and T follicular regulatory cells have different TCR specificity. Nat. Commun. 8, 15067. 10.1038/ncomms15067

35. Mamedov, I.Z., Britanova, O.V., Zvyagin, I.V., Turchaninova, M.A., Bolotin, D.A., Putintseva, E.V., Lebedev, Y.B., Chudakov, D.M., 2013. Preparing Unbiased T-Cell Receptor and Antibody cDNA Libraries for the Deep Next Generation Sequencing Profiling. Front. Immunol. 4. 10.3389/fimmu.2013.00456

36. Manfras, B.J., Terjung, D., Boehm, B.O., 1999. Non-productive human TCR β chain genes represent V-D-J diversity before selection upon function: insight into biased usage of TCRBD and TCRBJ genes and diversity of CDR3 region length. Hum. Immunol. 60, 1090–1100. 10.1016/S0198-8859(99)00099-3

37. Meynard-Piganeau, B., Feinauer, C., Weigt, M., Walczak, A.M., Mora, T., 2024. TULIP: A transformer-based unsupervised language model for interacting peptides and T cell receptors that generalizes to unseen epitopes. Proc. Natl. Acad. Sci. 121, e2316401121. 10.1073/pnas.2316401121

38. Minervina, A.A., Komech, E.A., Titov, A., Bensouda Koraichi, M., Rosati, E., Mamedov, I.Z., Franke, A., Efimov, G.A., Chudakov, D.M., Mora, T., Walczak, A.M., Lebedev, Y.B., Pogorelyy, M.V., 2021. Longitudinal high-throughput TCR repertoire profiling reveals the dynamics of T-cell memory formation after mild COVID-19 infection. eLife 10, e63502. 10.7554/eLife.63502

39. Mitchell, A.M., Baschal, E.E., McDaniel, K.A., Fleury, T., Choi, H., Pyle, L., Yu, L., Rewers, M.J., Nakayama, M., Michels, A.W., 2023. Tracking DNA-based antigen-specific T cell receptors during progression to type 1 diabetes. Sci. Adv. 9, eadj6975. 10.1126/sciadv.adj6975

40. Mora, T., Walczak, A., 2016. Quantifying lymphocyte receptor diversity.

41. Moris, P., De Pauw, J., Postovskaya, A., Gielis, S., De Neuter, N., Bittremieux, W., Ogunjimi, B., Laukens, K., Meysman, P., 2021. Current challenges for unseen-epitope TCR interaction prediction and a new perspective derived from image classification. Brief. Bioinform. 22, bbaa318. 10.1093/bib/bbaa318

42. Nagano, Y., Chain, B., 2023. tidytcells: standardizer for TR/MH nomenclature. Front. Immunol. 14. 10.3389/fimmu.2023.1276106

43. Nielsen, M., Eugster, A., Jensen, M.F., Goel, M., Tiffeau-Mayer, A., Pelissier, A., Valkiers, S., Martínez, M.R., Meynard-Piganeeau, B., Greiff, V., Mora, T., Walczak, A.M., Croce, G., Moreno, D.L., Gfeller, D., Meysman, P., Barton, J., 2024. Lessons learned from the IMMREP23 TCR-epitope prediction challenge. ImmunoInformatics 16, 100045. 10.1016/j.immuno.2024.100045

44. Nikolich-Žugich, J., Slifka, M.K., Messaoudi, I., 2004. The many important facets of T-cell repertoire diversity. Nat. Rev. Immunol. 4, 123–132. 10.1038/nri1292

45. Nishio, J., Suzuki, M., Nanki, T., Miyasaka, N., Kohsaka, H., 2004. Development of TCRB CDR3 length repertoire of human T lymphocytes. Int. Immunol. 16, 423–431. 10.1093/intimm/dxh046

46. Nolan, S., Vignali, M., Klinger, M., Dines, J.N., Kaplan, I.M., Svejnoha, E., Craft, T., Boland, K., Pesesky, M.W., Gittelman, R.M., Snyder, T.M., Gooley, C.J., Semprini, S., Cerchione, C., Nicolini, F., Mazza, M., Delmonte, O.M., Dobbs, K., Carreño-Tarragona, G., Barrio, S., Sambri, V., Martinelli, G., Goldman, J.D., Heath, J.R., Notarangelo, L.D., Martinez-Lopez, J., Howie, B., Carlson, J.M., Robins, H.S., 2025. A large-scale database of T-cell receptor beta sequences and binding associations from natural and synthetic exposure to SARS-CoV-2. Front. Immunol. 16. 10.3389/fimmu.2025.1488851

47. Omer, A., Peres, A., Rodriguez, O.L., Watson, C.T., Lees, W., Polak, P., Collins, A.M., Yaari, G., 2022. T cell receptor beta germline variability is revealed by inference from repertoire data. Genome Med. 14, 2. 10.1186/s13073-021-01008-4

48. Pannetier, C., Cochet, M., Darche, S., Casrouge, A., Zöller, M., Kourilsky, P., 1993. The sizes of the CDR3 hypervariable regions of the murine T-cell receptor beta chains vary as a function of the recombined germ-line segments. Proc. Natl. Acad. Sci. U. S. A. 90, 4319–4323. 10.1073/pnas.90.9.4319

49. Pham, M.-D.N., Nguyen, T.-N., Tran, L.S., Nguyen, Q.-T.B., Nguyen, T.-P.H., Pham, T.M.Q., Nguyen, H.-N., Giang, H., Phan, M.-D., Nguyen, V., 2023. epiTCR: a highly sensitive predictor for TCR–peptide binding. Bioinformatics 39, btad284. 10.1093/bioinformatics/btad284

50. Qi, Q., Liu, Y., Cheng, Y., Glanville, J., Zhang, D., Lee, J.-Y., Olshen, R.A., Weyand, C.M., Boyd, S.D., Goronzy, J.J., 2014. Diversity and clonal selection in the human T-cell repertoire. Proc. Natl. Acad. Sci. 111, 13139–13144. 10.1073/pnas.1409155111

51. Robins, H.S., Campregher, P.V., Srivastava, S.K., Wacher, A., Turtle, C.J., Kahsai, O., Riddell, S.R., Warren, E.H., Carlson, C.S., 2009. Comprehensive assessment of T-cell receptor β-chain diversity in αβ T cells. Blood 114, 4099–4107. 10.1182/blood-2009-04-217604

52. Rosenberg, S.A., Yang, J.C., Sherry, R.M., Kammula, U.S., Hughes, M.S., Phan, G.Q., Citrin, D.E., Restifo, N.P., Robbins, P.F., Wunderlich, J.R., Morton, K.E., Laurencot, C.M., Steinberg, S.M., White, D.E., Dudley, M.E., 2011. Durable complete responses in heavily pretreated patients with metastatic melanoma using T-cell transfer immunotherapy. Clin. Cancer Res. Off. J. Am. Assoc. Cancer Res. 17, 4550–4557. 10.1158/1078-0432.CCR-11-0116

53. Rubelt, F., Bolen, C.R., McGuire, H.M., Heiden, J.A.V., Gadala-Maria, D., Levin, M., M. Euskirchen, G., Mamedov, M.R., Swan, G.E., Dekker, C.L., Cowell, L.G., Kleinstein, S.H., Davis, M.M., 2016. Individual heritable differences result in unique cell lymphocyte receptor repertoires of naïve and antigen-experienced cells. Nat. Commun. 7, 11112. 10.1038/ncomms11112

54. Rudqvist, N.-P., Pilones, K.A., Lhuillier, C., Wennerberg, E., Sidhom, J.-W., Emerson, R.O., Robins, H.S., Schneck, J., Formenti, S.C., Demaria, S., 2018. Radiotherapy and CTLA-4 Blockade Shape the TCR Repertoire of Tumor-Infiltrating T Cells. Cancer Immunol. Res. 6, 139–150. 10.1158/2326-6066.CIR-17-0134

55. Shomuradova, A.S., Vagida, M.S., Sheetikov, S.A., Zornikova, K.V., Kiryukhin, D., Titov, A., Peshkova, I.O., Khmelevskaya, A., Dianov, D.V., Malasheva, M., Shmelev, A., Serdyuk, Y., Bagaev, D.V., Pivnyuk, A., Shcherbinin, D.S., Maleeva, A.V., Shakirova, N.T., Pilunov, A., Malko, D.B., Khamaganova, E.G., Biderman, B., Ivanov, A., Shugay, M., Efimov, G.A., 2020. SARS-CoV-2 Epitopes Are Recognized by a Public and Diverse Repertoire of Human T Cell Receptors. Immunity 53, 1245–1257.e5. 10.1016/j.immuni.2020.11.004

56. Shugay, M., Bagaev, D.V., Turchaninova, M.A., Bolotin, D.A., Britanova, O.V., Putintseva, E.V., Pogorelyy, M.V., Nazarov, V.I., Zvyagin, I.V., Kirgizova, V.I., Kirgizov, K.I., Skorobogatova, E.V., Chudakov, D.M., 2015. VDJtools: Unifying Post-analysis of T Cell Receptor Repertoires. PLOS Comput. Biol. 11, e1004503. 10.1371/journal.pcbi.1004503

57. Shugay, M., Bagaev, D.V., Zvyagin, I.V., Vroomans, R.M., Crawford, J.C., Dolton, G., Komech, E.A., Sycheva, A.L., Koneva, A.E., Egorov, E.S., Eliseev, A.V., Van Dyk, E., Dash, P., Attaf, M., Rius, C., Ladell, K., McLaren, J.E., Matthews, K.K., Clemens, E.B., Douek, D.C., Luciani, F., van Baarle, D., Kedzierska, K., Kesmir, C., Thomas, P.G., Price, D.A., Sewell, A.K., Chudakov, D.M., 2018. VDJdb: a curated database of T-cell receptor sequences with known antigen specificity. Nucleic Acids Res. 46, D419–D427. 10.1093/nar/gkx760

58. Shugay, M., Britanova, O.V., Merzlyak, E.M., Turchaninova, M.A., Mamedov, I.Z., Tuganbaev, T.R., Bolotin, D.A., Staroverov, D.B., Putintseva, E.V., Plevova, K., Linnemann, C., Shagin, D., Pospisilova, S., Lukyanov, S., Schumacher, T.N., Chudakov, D.M., 2014. Towards error-free profiling of immune repertoires. Nat. Methods 11, 653–655. 10.1038/nmeth.2960

59. Sims, J.S., Grinshpun, B., Feng, Y., Ung, T.H., Neira, J.A., Samanamud, J.L., Canoll, P., Shen, Y., Sims, P.A., Bruce, J.N., 2016. Diversity and divergence of the glioma-infiltrating T-cell receptor repertoire. Proc. Natl. Acad. Sci. U. S. A. 113, E3529–3537. 10.1073/pnas.1601012113

60. Theil, A., Wilhelm, C., Kuhn, M., Petzold, A., Tuve, S., Oelschlägel, U., Dahl, A., Bornhäuser, M., Bonifacio, E., Eugster, A., 2017. T cell receptor repertoires after adoptive transfer of expanded allogeneic regulatory T cells. Clin. Exp. Immunol. 187, 316–324. 10.1111/cei.12887

61. Tickotsky, N., Sagiv, T., Prilusky, J., Shifrut, E., Friedman, N., 2017. McPAS-TCR: a manually curated catalogue of pathology-associated T cell receptor sequences. Bioinformatics 33, 2924–2929. 10.1093/bioinformatics/btx286

62. Valkiers, S., Van Houcke, M., Laukens, K., Meysman, P., 2021. ClusTCR: a python interface for rapid clustering of large sets of CDR3 sequences with unknown antigen specificity. Bioinformatics 37, 4865–4867. 10.1093/bioinformatics/btab446

63. Venturi, V., Nzingha, K., Amos, T.G., Charles, W.C., Dekhtiarenko, I., Cicin-Sain, L., Davenport, M.P., Rudd, B.D., 2016. The Neonatal CD8+ T Cell Repertoire Rapidly Diversifies during Persistent Viral Infection. J. Immunol. 196, 1604–1616. 10.4049/jimmunol.1501867

64. Vita, R., Mahajan, S., Overton, J.A., Dhanda, S.K., Martini, S., Cantrell, J.R., Wheeler, D.K., Sette, A., Peters, B., 2019. The Immune Epitope Database (IEDB): 2018 update. Nucleic Acids Res. 47, D339–D343. 10.1093/nar/gky1006

65. Vita, R., Overton, J.A., Greenbaum, J.A., Ponomarenko, J., Clark, J.D., Cantrell, J.R., Wheeler, D.K., Gabbard, J.L., Hix, D., Sette, A., Peters, B., 2015. The immune epitope database (IEDB) 3.0. Nucleic Acids Res. 43, D405–D412. 10.1093/nar/gku938

66. Westcott, P.M.K., Sacks, N.J., Schenkel, J.M., Ely, Z.A., Smith, O., Hauck, H., Jaeger, A.M., Zhang, D., Backlund, C.M., Beytagh, M.C., Patten, J.J., Elbashir, R., Eng, G., Irvine, D.J., Yilmaz, O.H., Jacks, T., 2021. Low neoantigen expression and poor T-cell priming underlie early immune escape in colorectal cancer. Nat. Cancer 2, 1071–1085. 10.1038/s43018-021-00247-z

67. Xu, A.M., Chour, W., DeLucia, D.C., Su, Y., Pavlovitch-Bedzyk, A.J., Ng, R., Rasheed, Y., Davis, M.M., Lee, J.K., Heath, J.R., 2023. Entropic analysis of antigen-specific CDR3 domains identifies essential binding motifs shared by CDR3s with different antigen specificities. Cell Syst. 14, 273–284.e5. 10.1016/j.cels.2023.03.001

68. Ye, J., Ma, N., Madden, T.L., Ostell, J.M., 2013. IgBLAST: an immunoglobulin variable domain sequence analysis tool. Nucleic Acids Res. 41, W34–40. 10.1093/nar/gkt382

69. Zappasodi, R., Sirard, C., Li, Y., Budhu, S., Abu-Akeel, M., Liu, C., Yang, X., Zhong, H., Newman, W., Qi, J., Wong, P., Schaer, D., Koon, H., Velcheti, V., Hellmann, M.D., Postow, M.A., Callahan, M.K., Wolchok, J.D., Merghoub, T., 2019. Rational design of anti-GITR-based combination immunotherapy. Nat. Med. 25, 759–766. 10.1038/s41591-019-0420-8

70. Zarnitsyna, V., Evavold, B., Schoettle, L., Blattman, J., Antia, R., 2013. Estimating the Diversity, Completeness, and Cross-Reactivity of the T Cell Repertoire. Front. Immunol. 4. 10.3389/fimmu.2013.00485

71. Zheng, F., Xu, H., Zhang, C., Hong, X., Liu, D., Tang, D., Xiong, Z., Dai, Y., 2021. Immune cell and TCR/BCR repertoire profiling in systemic lupus erythematosus patients by single-cell sequencing. Aging 13, 24432–24448. 10.18632/aging.203695

