## Supplementary Tables and Figures for "Statistical modelling of CDR3 sequences provides robust quality control for TCR repertoire datasets"

**The PDF file includes:**

Tables S1 to S3

Figs. S1 to S6

### Supplementary Tables

**Supplementary Table 1.** List of iReceptor studies for the human TCR repertoire datasets considered in this work (*TRA and TRB data were available for the study PRJNA300878 on the iReceptor website even if TRA is not indicated in the interface).

| Reference | iReceptor study ID | Total number of TCRs | Number of subjects | Chain |
|---|---|---|---|---|
| (Shomuradova et al., 2020) | IR-Efimov-000001 | 133,706 | 11 | TRA |
| (Shomuradova et al., 2020) | IR-Efimov-000002 | 23,720 | 2 | TRA |
| (Rubelt et al., 2016) | PRJNA300878* | 92,588 | 10 | TRA |
| (Minervina et al., 2021) | PRJNA633317 | 6,056,676 | 2 | TRA |
| (Hannan et al., 2021) | 4505707319090933270-242ac113-0001-012 | 580,620 | 8 | TRB |
| (Shomuradova et al., 2020) | IR-Efimov-000001 | 810,514 | 12 | TRB |
| (Shomuradova et al., 2020) | IR-Efimov-000002 | 1,661,054 | 16 | TRB |
| (Joseph et al., 2022) | IR-Roche-000001 | 5,151,261 | 95 | TRB |
| (Nolan et al., 2025) | ImmuneCODE-COVID-Release-002: COVID-19-BWNW | 15,579,052 | 50 | TRB |
| (Nolan et al., 2025) | ImmuneCODE-COVID-Release-002: COVID-19-HUniv12Oc | 33,007,109 | 170 | TRB |
| (Nolan et al., 2025) | ImmuneCODE-COVID-Release-002: COVID-19-IRST/AUSL | 9,682,107 | 64 | TRB |
| (Nolan et al., 2025) | ImmuneCODE-COVID-Release-002: COVID-19-ISB | 33,865,273 | 83 | TRB |
| (Nolan et al., 2025) | ImmuneCODE-COVID-Release-002: COVID-19-NIH/NIAID | 25,584,351 | 299 | TRB |
| (Rubelt et al., 2016) | PRJNA300878 | 151,965 | 10 | TRB |
| (Jia et al., 2018) | PRJNA506151 | 441,296 | 15 | TRB |
| (Minervina et al., 2021) | PRJNA633317 | 13,783,943 | 2 | TRB |
| (DeWitt et al., 2015) | dewitt-2015-jvi | 5,059,222 | 9 | TRB |

**Supplementary Table 2.** List of repositories for the murine TCR repertoire dataset considered in this work.

| Reference | Total number of TCRs | Number of subjects | Chain | Source of data |
|---|---|---|---|---|
| (Brown et al., 2024) | 1,067,107 | 70 | TRA | https://zenodo.org/doi/10.5281/zenodo.11081248 |
| (Genolet et al., 2023) | 106,028 | 6 | TRA | GEO:GSE225984 |
| (Brown et al., 2021) | 64,581 | 18 | TRB | Adaptive Biotech |
| (Dudgeon et al., 2014) | 126,106 | 2 | TRB | Adaptive Biotech |
| adaptivebiotech.com/pub/mouse-skin-2016 | 2,995 | 3 | TRB | Adaptive Biotech |
| (Gaide et al., 2015) | 153,828 | 42 | TRB | Adaptive Biotech |
| (Genolet et al., 2023) | 224,756 | 6 | TRB | GEO:GSE225984 |
| adaptivebiotech.com/pub/2-mouse-strain-comparison | 1,018,166 | 8 | TRB | Adaptive Biotech |
| (Iijima and Iwasaki, 2014) | 1,484 | 2 | TRB | Adaptive Biotech |
| (Rudqvist et al., 2018) | 51,434 | 54 | TRB | Adaptive Biotech |
| (Venturi et al., 2016) | 154,413 | 5 | TRB | Adaptive Biotech |
| (Westcott et al., 2021) | 333 | 8 | TRB | Adaptive Biotech |
| (Zappasodi et al., 2019) | 144,572 | 6 | TRB | Adaptive Biotech |

**Supplementary Table 3.** List of iReceptor repositories with a high number of predicted inconsistencies used in Figure 4.

| | Reference | iReceptor study ID | Total number of TCRs | Number of subjects | Chain |
|---|---|---|---|---|---|
| Study 1 | (Lang Kuhs et al., 2018) | langkuhs-2018-plosone | 2,007,389 | 50 | TRB |
| Study 2 | (Theil et al., 2017) | 1371444213709729305-242ac11c-0001-012 | 4,260,102 | 4 | TRA |
| Study 3 | (Sims et al., 2016) | PRJNA315543 | 4,946,968 | 15 | TRA |
| Study 4 | (Maceiras et al., 2017) | PRJNA362309 | 119,935 | 4 | TRA |

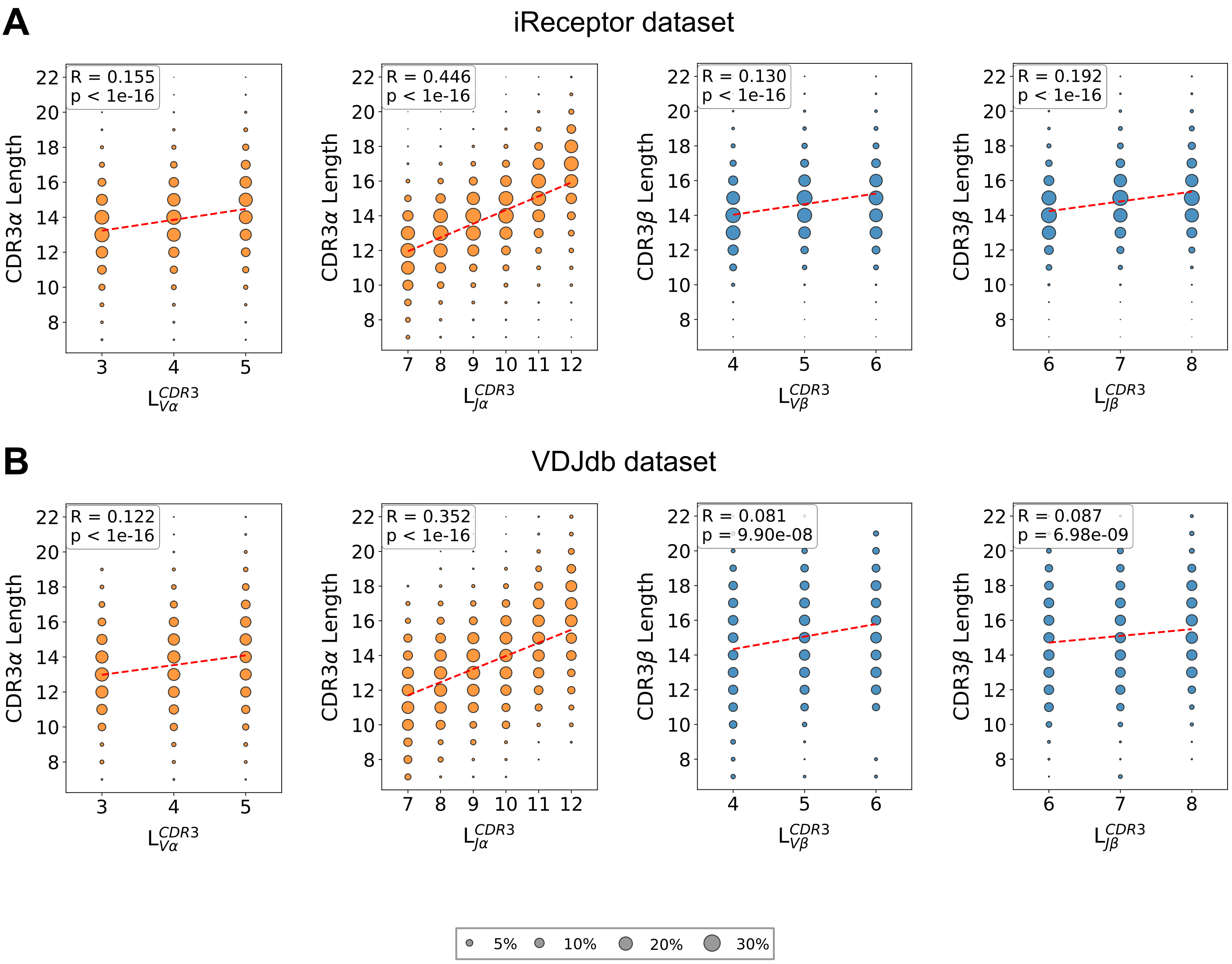


**Supplementary Figure 1**: V and J gene usage shapes the length of CDR3 sequences. **(A)** Correlations between CDR3 length and the number of CDR3 residues for each germline V or J gene ($L^{CDR3}{}_{V}$ or $L^{CDR3}{}_{J}$) for α and β chains in human TCR repertoire data. The size of each dot is proportional to the number of sequences. **(B)** Correlations between CDR3 length and the number of CDR3 residues for each germline V or J gene ($L^{CDR3}{}_{V}$ or $L^{CDR3}{}_{J}$) for α and β chains in human epitope-specific VDJdb dataset. The size of each dot is proportional to the number of sequences.

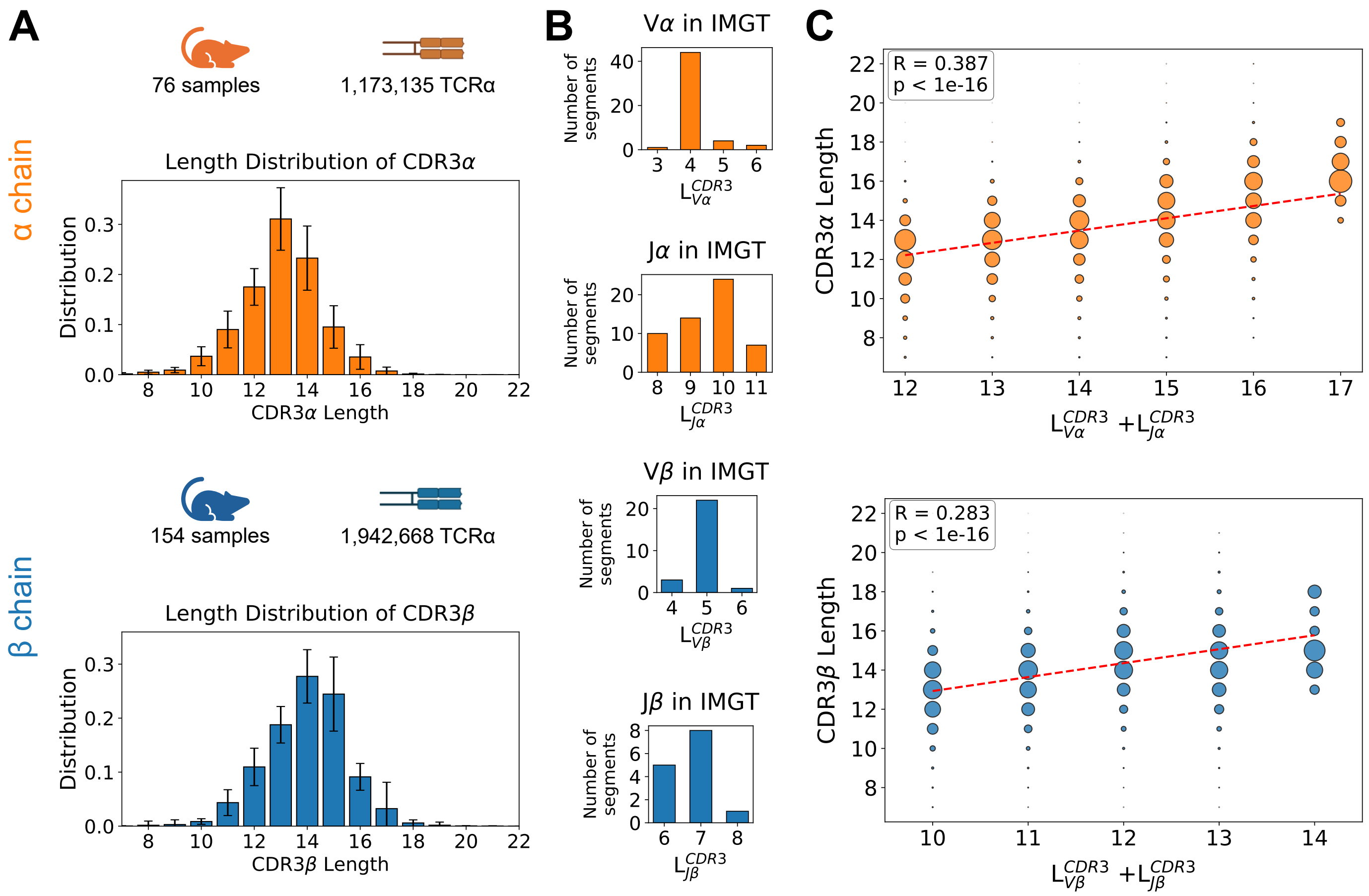


**Supplementary Figure 2:** V and J gene usage shapes the length of CDR3 sequences in murine data. **(A)** Summary of the murine TCR repertoire datasets and distributions of CDR3 lengths for α and β chains. Error bars show the standard deviation across subjects. **(B)** Distributions of the number of CDR3 residues in germline encoded V and J genes ($L^{CDR3}{}_V$ and $L^{CDR3}{}_J$) for α and β chains based on the IMGT reference database. **(C)** Correlations between the number of CDR3 residues in germline V and J genes ($L^{CDR3}{}_V$+$L^{CDR3}{}_J$) and CDR3 length for both chains. The size of each dot is proportional to the number of sequences.

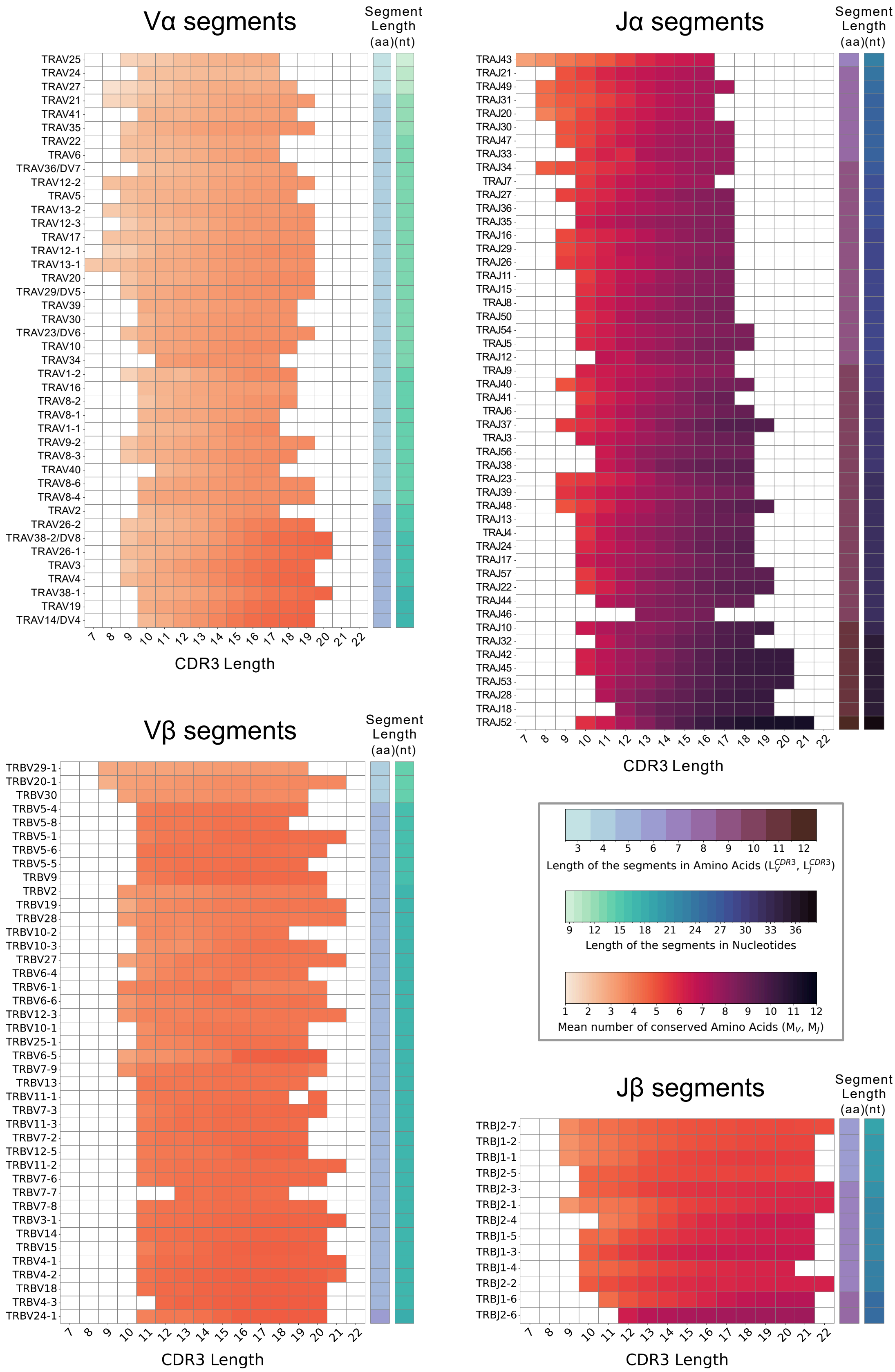


**Supplementary Figure 3:** Heatmaps showing $M_{V,L}$ and $M_{J,L}$ in human TCR repertoire data for Vα, Jα, Vβ and Jβ genes.

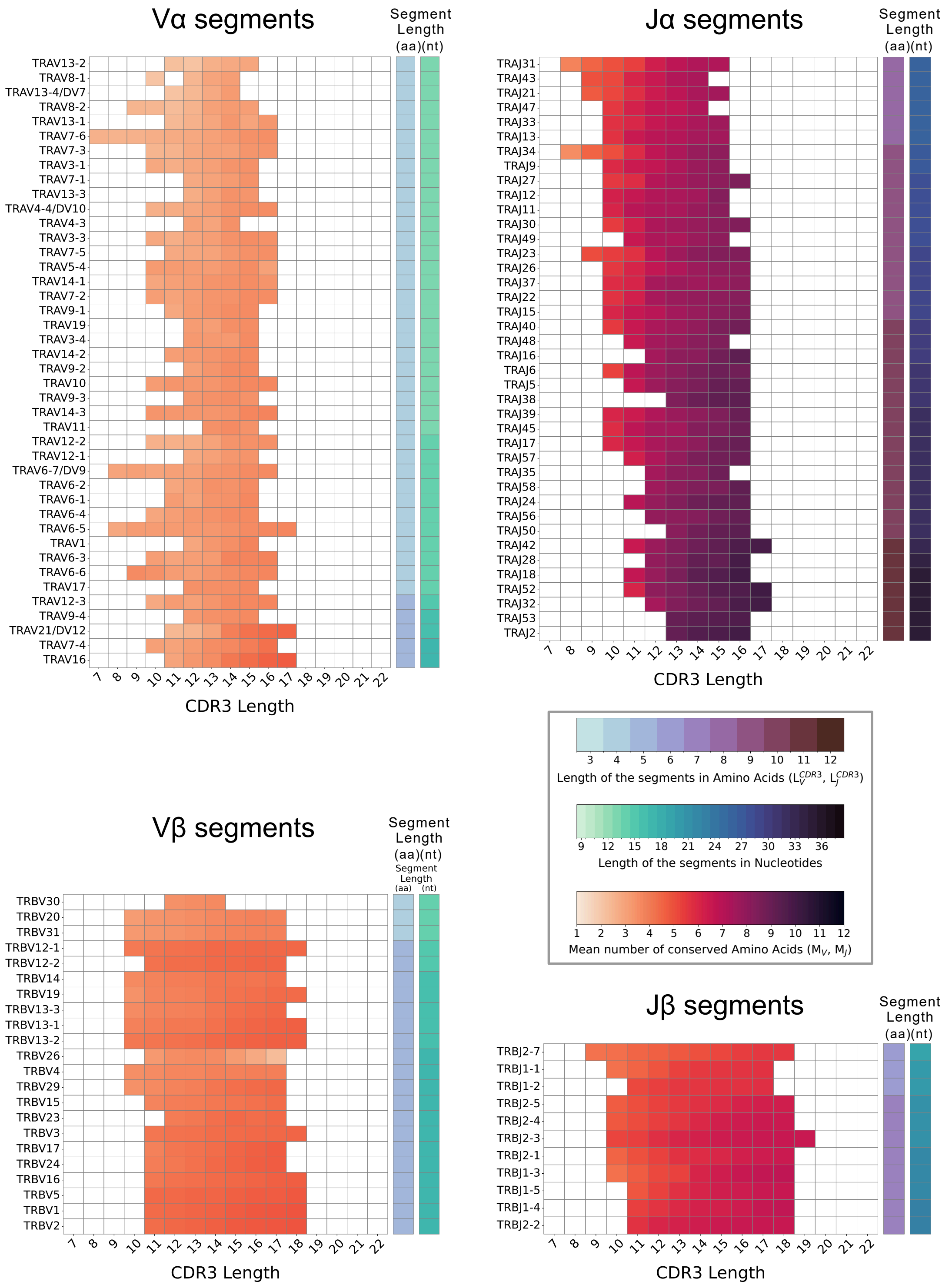


**Supplementary Figure 4:** Heatmaps showing $M_{V,L}$ and $M_{J,L}$ in murine TCR repertoire data for Vα, Jα, Vβ and Jβ genes.

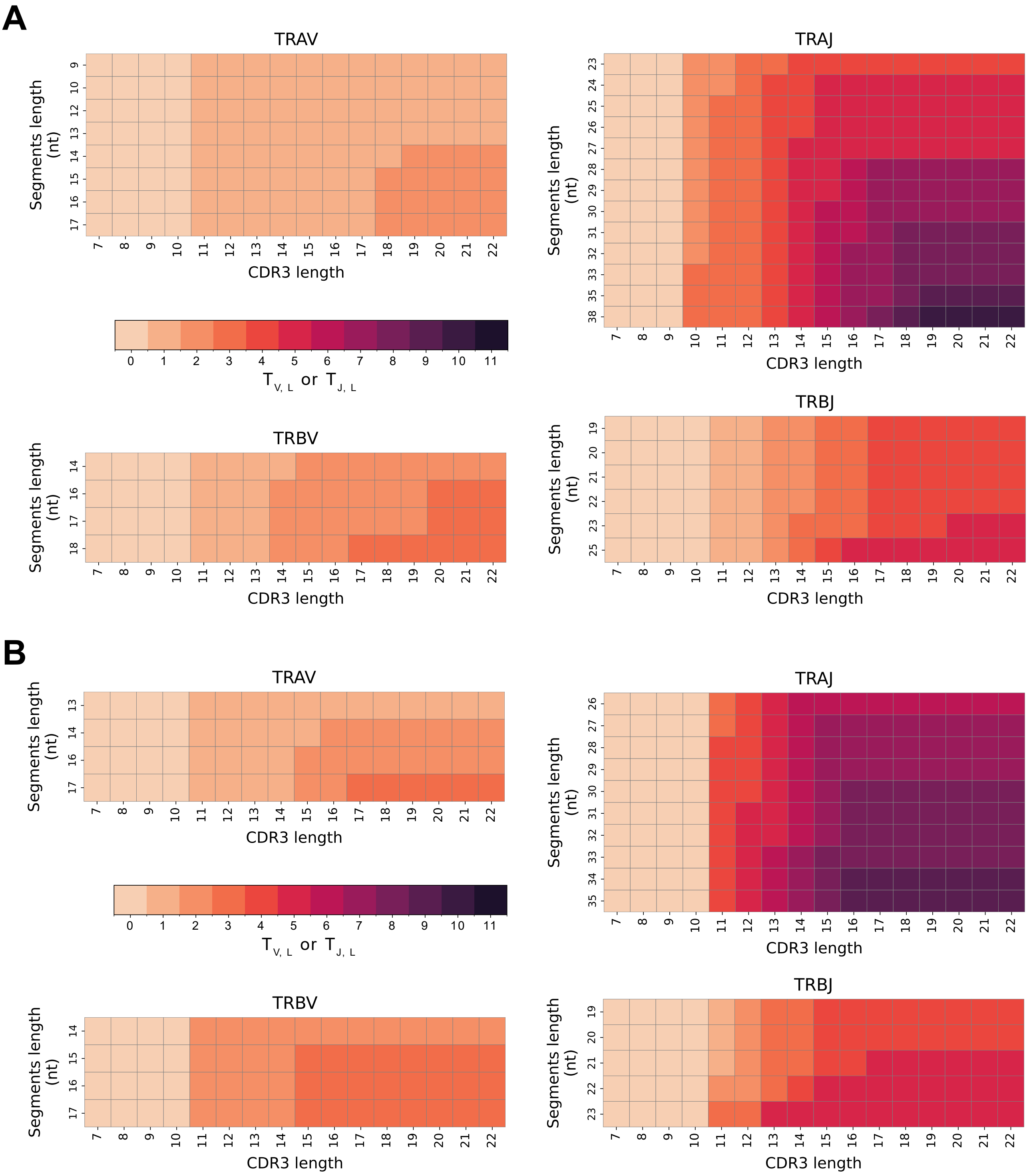


**Supplementary Figure 5:** Expected number of consistently conserved germline residues $T_{V,L}$ and $T_{J,L}$ derived for **(A)** human TCR repertoires and **(B)** murine TCR repertoires.

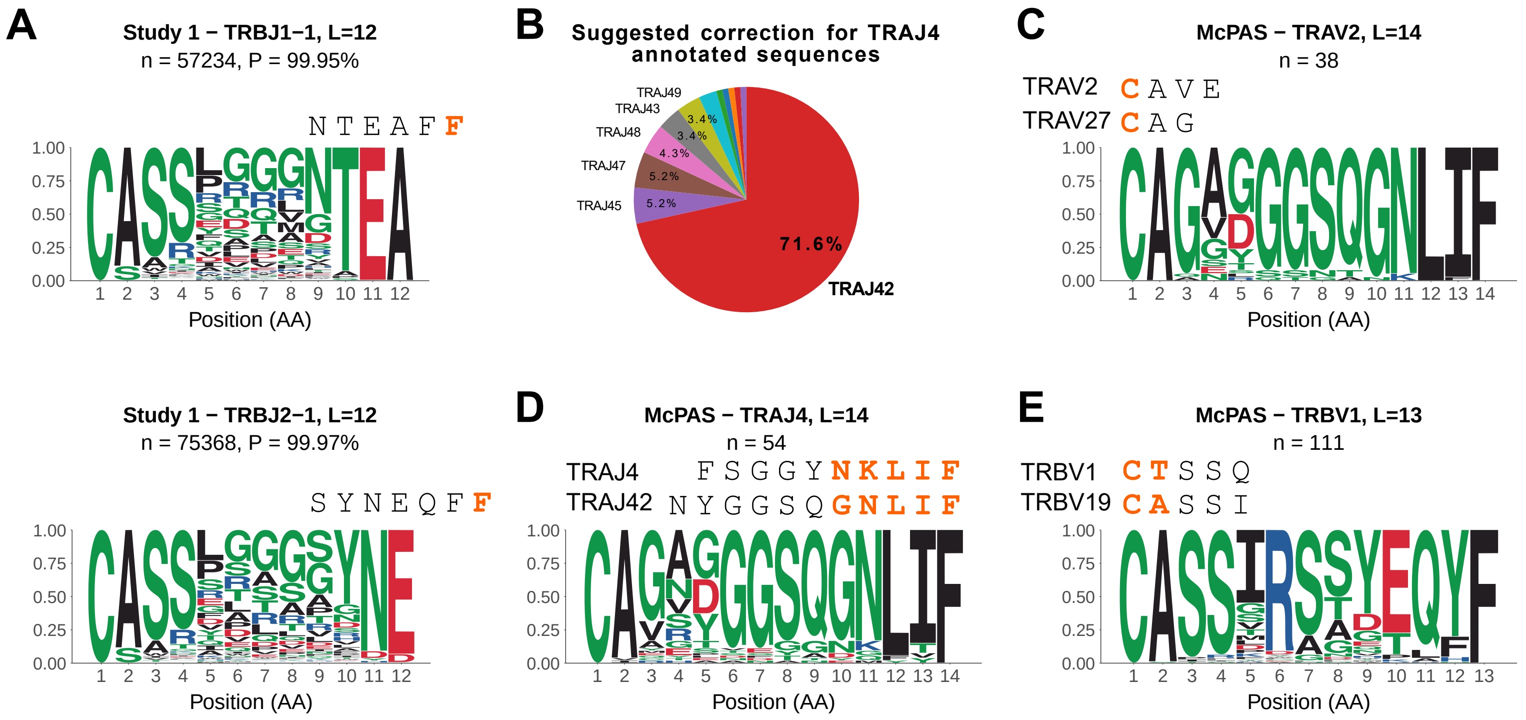


**Supplementary Figure 6:** Consistency between V and J gene usage and CDR3 sequences provides a robust quality control for TCR repertoire datasets. **(A)** Example of systematic truncations of two or three C-terminal residues in CDR3β sequences of TRBJ1-1 and TRBJ2-1 TCRs predicted as inconsistent (Study 1). **(B)** Pie chart showing the distribution of Jα genes inferred from CDR3 sequences for TCR sequences annotated as TRAJ4 in the McPAS GILGFVFTL-specific dataset. **(C)** Motifs of all CDR3α sequences of TRAV2 TCRs of length 14 in the McPAS GILGFVFTL-specific dataset showing better consistency with the TRAV27 germline sequence. **(D)** Motifs of all CDR3α sequences of TRAJ4 TCRs of length 14 in the McPAS GILGFVFTL-specific dataset showing better consistency with the TRAJ42 germline sequence. **(E)** Motifs of all CDR3β sequences of TRBV1 (pseudo-gene) TCRs of length 13 in the McPAS GILGFVFTL-specific dataset showing better consistency with TRBV19 germline sequence. 'P' stands for the proportion of predicted inconsistencies within each particular subset and 'n' for the number of sequences from which the motif was built. The orange letters represent the germline amino acids expected to be conserved based on $T_{V,\,L}$ and $T_{J,\,L}$ thresholds.
